# The Effects of Dual-Site Beta tACS over the rIFG and preSMA on Response Inhibition in Young and Older Adults

**DOI:** 10.1101/2022.11.30.518460

**Authors:** Jane Tan, Kartik K. Iyer, Michael A. Nitsche, Rohan Puri, Mark R. Hinder, Hakuei Fujiyama

**Author notes:** **Corresponding Author:** Jane Tan, Discipline of Psychology, College of Science, Health, Engineering and Education, Murdoch University, Perth, Australia, 90 South Street, Murdoch, WA, 6150, Australia.

## Abstract

A growing body of research suggests that changes in both structural and functional connectivity in the aging brain contribute to declines in cognitive functions such as response inhibition. In recent years, transcranial alternating current stimulation (tACS) has garnered substantial research interest as a potential tool for the modulation of functional connectivity. Here, we report the findings from a double-blind crossover study that investigated the effects of dual-site beta tACS over the right inferior frontal gyrus (rIFG) and pre-supplementary motor area (preSMA) on functional connectivity measured with electroencephalography and response inhibition (stop-signal task performance) of healthy young (*n* = 18, aged 18-34 years) and older (*n* =15, aged 61-79 years) adults. Two tACS conditions were administered in separate sessions: in-phase tACS, where electrical currents delivered to the rIFG and preSMA had a 0° phase difference, and anti-phase tACS, where currents had a 180° phase difference. Stop-signal task performance was assessed before and after tACS. We found significant improvements in response inhibition that were not due to the phase of the tACS applied. There were also no significant changes in rIFG-preSMA phase connectivity in either age group from in- or anti-phase tACS. Furthermore, we did not observe significant differences in rIFG-preSMA phase connectivity between successful and unsuccessful inhibition, which suggests that rIFG-preSMA phase-coupling might not underlie effective response inhibition. The results offer insight into the neurophysiology of response inhibition and contribute to the future development of non-pharmacological interventions aimed at alleviating age-related declines in cognitive function.

## 1. Introduction

Response inhibition, or the ability to cancel a pre-potent motor action, is a fundamental cognitive function that is crucial for flexible adaptation to a changing external environment (Miyake et al., 2000). However, response inhibition tends to decline as we age (e.g., Hermans et al., 2018), with deleterious effects on the ability of older adults to engage in daily activities safely and independently. Indeed, poorer response inhibition has been linked to a higher risk of falls for older individuals (Schoene et al., 2017). The development of effective and reliable interventions to alleviate age-related inhibitory deficits could therefore play a critical role in improving the well-being of elderly populations.

There is mounting evidence suggesting that age-related changes in the structural and functional connections within brain networks contribute to age-related cognitive decline. Greater declines in executive functioning and attention have been associated with lower white matter structural integrity and decreased segregation of brain networks such as the default mode and executive control networks (Brown et al., 2019; Chong et al., 2019). Notably, the functional connectivity between the right inferior frontal gyrus (rIFG), pre-supplementary motor area (preSMA), and sensorimotor regions has been found to play a progressively larger role in response inhibition performance in older individuals (Tsvetanov et al., 2018). It has been postulated that functional interactions between the rIFG and preSMA subserve connectivity within the fronto-basal-ganglia network, which underlies response inhibition performance (Xu et al., 2016). As outlined in a previous review (Tan et al., 2019), we proposed that facilitating the functional connectivity between the rIFG and the preSMA, by way of non-invasive brain stimulation, may mitigate age-related inhibitory deficits.

Transcranial alternating current stimulation (tACS) – a form of non-invasive brain stimulation – has garnered substantial research interest as a potential tool for the modulation of functional connectivity. This technique involves the application of weak alternating electrical currents through electrodes placed over the scalp and has been primarily used to modulate neural oscillatory activity (Antal & Herrmann, 2016). While in-depth understanding of tACS mechanisms is still being advanced by emerging research, there is evidence to support the frequency-specific entrainment of neural activity during tACS application. For instance, the phase-locking of endogenous neural activity to the stimulation frequency was found to be significantly stronger during real tACS when compared with sham stimulation (Helfrich, Schneider, et al., 2014; Wischnewski, Engelhardt, et al., 2019). Single-neuron recordings in non-human primates during tACS administration indicated that spike timings occurred preferentially around the 0° phase of each tACS cycle, and exhibited higher phase-locking values when compared with sham stimulation (Krause et al., 2019).

Based on the concept of entrainment, tACS has been shown to exert modulatory effects on oscillatory phasic relationships *between* distinct cortical regions. Through the simultaneous recording of electroencephalographic (EEG) activity during tACS application, the delivery of alternating currents at the same phase (in-phase, i.e., with 0° phase difference) to two brain regions (i.e., dual-site) was shown to strengthen the frequency-specific phase-coupling of electrophysiological signals between those stimulated sites (Helfrich, Knepper, et al., 2014; Schwab et al., 2019). The effects of dual-site tACS have also been found to be phase-dependent, such that the phase-synchronisation of two regions was weakened when tACS currents were delivered at a 180° phase difference (i.e., anti-phase tACS) (Helfrich, Knepper, et al., 2014). The increase in phase-coupling elicited by in-phase tACS was also significantly larger when compared with that of jittered-phase and sham stimulation (Helfrich, Knepper, et al., 2014; Schwab et al., 2019). These findings support the communication-through-coherence theory, which posits that the strength of inter- and intra-regional connectivity in the brain is underpinned by the oscillatory phase relationships of neuronal groups, with higher phase coherence facilitating neuronal information transfer and functional connectivity (Fries, 2005, 2015; Womelsdorf et al., 2007). Furthermore, phase-coherent neural oscillations are thought to be spectral signatures of the large-scale/inter-regional interactions between and within neural networks that underlie cognitive processing (Salinas & Sejnowski, 2001; Siegel et al., 2012). Crucially, Polanía et al. (2012) found evidence suggesting that inter-regional phase-coupling is *causally* involved in cognitive functioning. Through the application of dual-site theta tACS to left fronto-parietal sites, the authors observed that in-phase tACS resulted in significantly improved verbal working memory when compared with sham and anti-phase stimulation (Polanía et al., 2012). Similarly, the delivery of in-phase gamma tACS to interhemispheric parietal-occipital regions was shown to improve motion perception compared to anti-phase stimulation, a likely consequence of the stronger functional coupling between the stimulated regions that resulted from in-phase tACS (Helfrich, Knepper, et al., 2014).

In light of the mounting evidence of strong associations between age-associated changes in functional networks and cognitive functioning, it was postulated that the facilitatory effects of in-phase tACS on functional connectivity can be harnessed for the alleviation of age-related cognitive declines (Tan et al., 2019). Indeed, this hypothesis was corroborated by Reinhart and Nguyen’s (2019) study, which found that in-phase tACS over fronto-temporal regions significantly improved working memory performance of older adults, with stronger theta phase-coupling between the stimulated regions. To the best of our knowledge, their findings constitute the first empirical evidence that this conceptual framework can be applied to the aging brain for the improvement of cognitive functions. The main objective of the current study was to examine the viability of this approach within the context of response inhibition. Namely, to determine if a dual-site tACS protocol that is designed to facilitate neural oscillatory phase-coupling can strengthen the functional connectivity between rIFG and preSMA and improve response inhibition in both young and older adults (Tan et al., 2019).

Using a double-blind crossover design, the current study investigated the effects of dual-site beta tACS over the rIFG and preSMA on response inhibition in young and older adults; changes in functional connectivity before and after stimulation were assessed using a series of EEG measures. Beta (20 Hz) was chosen as the stimulation frequency in view of its putative role in information transfer within the fronto-basal-ganglia network during response inhibition (Aron et al., 2016; Swann et al., 2011). Our primary hypothesis was that dual-site tACS would modulate response inhibition and resting-state functional connectivity in a phase-dependent and frequency-specific manner: in-phase stimulation was expected to facilitate beta-band phase-coupling between the stimulated regions and improve response inhibition of participants in both age groups. Conversely, anti-phase tACS was anticipated to degrade response inhibition and weaken the phase connectivity between the rIFG and preSMA.

## 2. Materials and Methods

### 2.1. Design

Participants underwent two stimulation conditions (in- and anti-phase tACS) that were administered in separate sessions (Figure 1), counterbalanced across participants, and conducted at least a week apart. Both sessions were conducted at the same time of day to control for any possible time-of-day cortisol level fluctuations that might affect tACS response (Sale et al., 2008). Resting-state (eyes-open and eyes-closed; 3 min each) and task-related EEG activity were recorded before and after the administration of tACS. However, only eyes-opened activity was analyzed as some participants reported having fallen asleep during the recording of eyes-closed activity. Furthermore, analysis of eyes-open resting-state activity is recommended for experimental designs utilising tasks that require visual processing (Barry et al., 2007), such as the stop-signal task in the current study. Task-related activity was measured during performance of the stop-signal task (described below).

**Figure 1.**
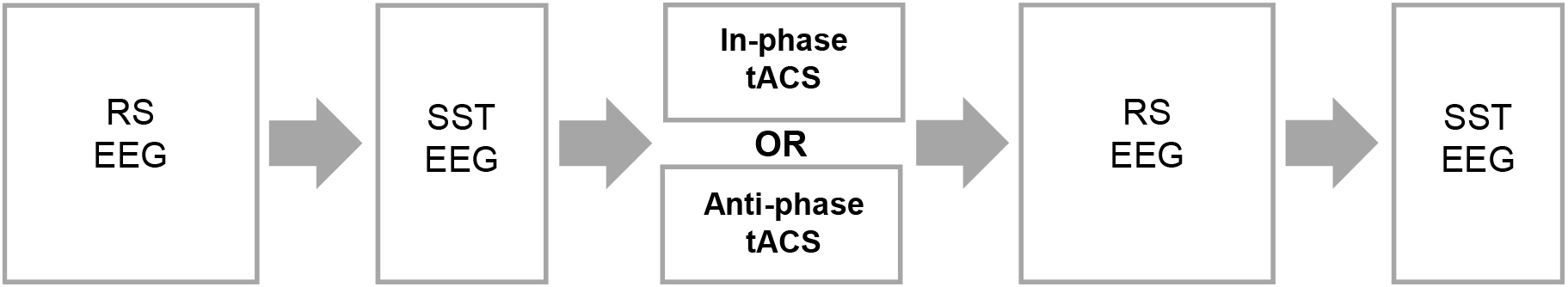
Experimental Protocol. *Note*. Resting-state (RS) and task-related EEG activity were measured before and after tACS. Task-related activity was recorded during stop-signal task (SST) performance. The two tACS conditions, in- and anti-phase, were administered in separate sessions.

### 2.2. Participants

Power analysis (power = 0.80; α = .05) indicated that 12 participants for each age group were required for an expected medium effect size of improved response inhibition by in-phase tACS. This effect size was based on the findings of previous studies that had investigated the effects of transcranial electrical stimulation on response inhibition (e.g., Hogeveen et al., 2016). In consideration of possible attrition, excessively noisy EEG recordings, and unsatisfactory task performance (see Section 2.6.1; Verbruggen et al., 2019), a total of 41 participants (20 young and 18 older adults) were recruited. The data of 5 participants were excluded from all analyses: two (1 young, 1 older) were excluded due to excessive movement artifacts in their EEG recordings, and three (1 young, 2 older) were excluded because of their stop-signal task performance (failed stop probability was < 0.25 or > 0.75). The final dataset comprised of 33 participants, with 18 young participants (11 females; mean age = 23.56 years, *SD* = 4.49 years) and 15 older participants (10 females; mean age = 68.80 years, *SD* = 5.39 years).

All participants were right-handed according to the Edinburgh Handedness Inventory (Oldfield, 1971; young participants: Laterality Quotient, LQ = 83.33 ± 15.34; older participants: LQ = 84.67 ± 18.85)). The Montreal Cognitive Assessment (MoCA; Nasreddine et al., 2012) was used to screen older participants for possible cognitive deficits, and all of them met Carson et al.’s (2018) recommended cut-off score of 23 (mean ± SD = 28.33 ± 1.40), suggesting no deficits. None of the participants reported any existing neurological conditions, metal implants or implanted neurostimulators, cochlear implants, cardiac pacemaker, and intra-cardiac lines. Apart from one older individual who was on a non-steroidal aromatase inhibitor, none of the participants were on psychoactive medications.

To assess possible confounds to behavior and neurophysiology, participants were asked during each session to report their quality of sleep on a rating scale (1: extremely poor sleep to 10: extremely good sleep) and number of hours slept during the previous night, and the amount of caffeine and alcohol they had consumed within 12 hours prior to the experimental session. A questionnaire was also administered to assess the occurrence and intensity of sensations (e.g., itching and tingling) that participants might have experienced while receiving tACS (Fertonani et al., 2015).

This study was approved by the Murdoch University Human Research Ethics Committee (2016/021). Participants provided written informed consent before taking part in the experiment and were remunerated with either academic credits or cash payments of $20 per session. This study was conducted in accordance with The Declaration of Helsinki (World Medical Association, 2013).

### 2.3. Stop-Signal Task

Response inhibition performance was assessed using a stop-signal task, whereby inhibitory performance was operationalized as the stop-signal reaction time (SSRT). The SSRT estimates the amount of time needed to cancel an ongoing motor response upon presentation of a stop signal (Logan & Cowan, 1984). The task was programmed using PsychoPy (version 1.83.04; Peirce et al., 2019) and displayed on a computer monitor with a refresh rate of 144 Hz.

Each trial began with the presentation of a fixation cross for a time interval that was jittered between 250 and 750 ms (drawn from a uniform distribution) before presentation of an imperative (go) signal, which was a green circle. Participants were asked to respond as quickly as possible by pressing the space bar on a computer keyboard with their right index finger whenever the imperative signal was presented (Figure 2).

**Figure 2.**
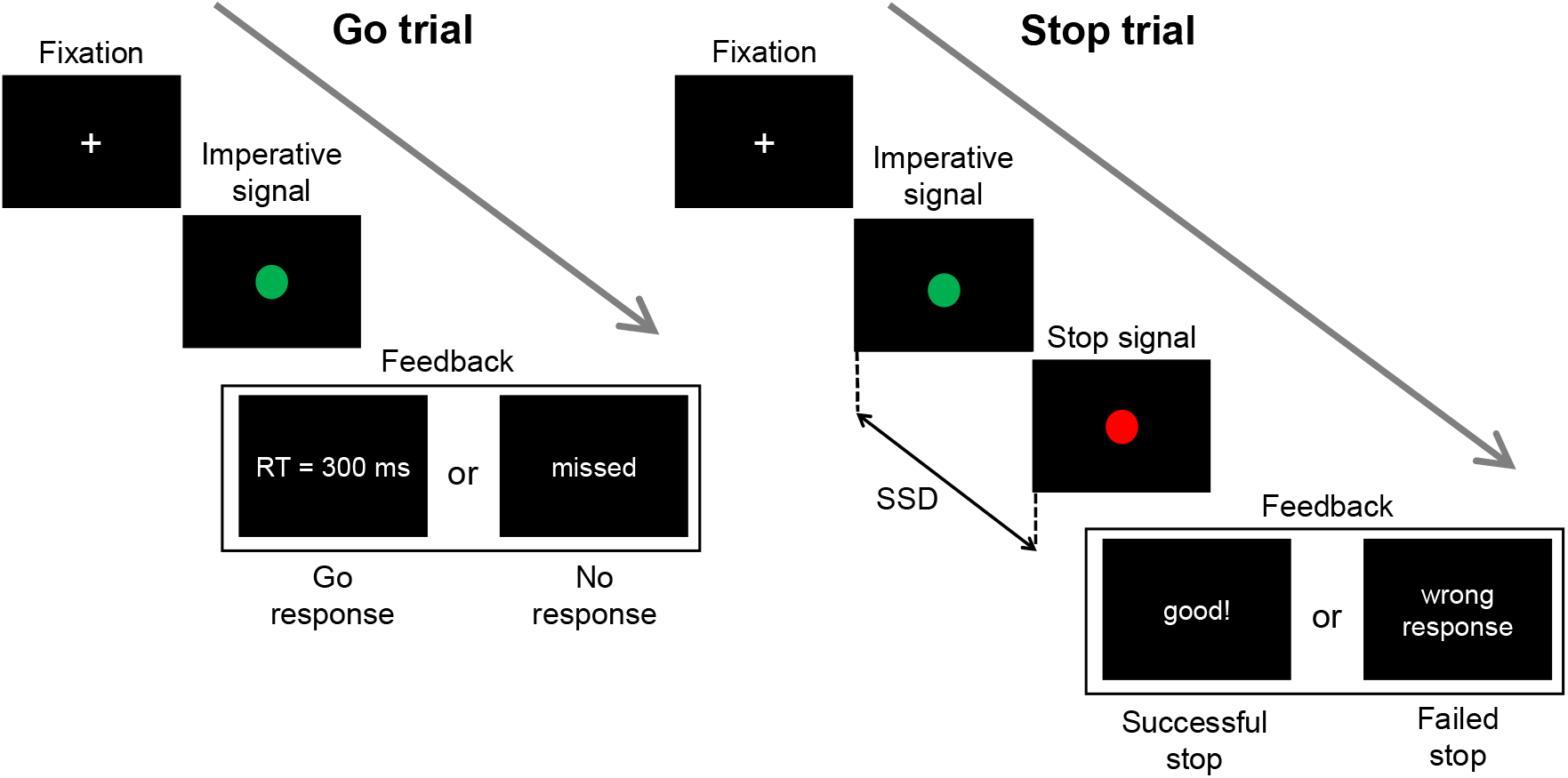
The Stop-Signal Task. *Note*. The imperative signal was a green circle, to which participants were asked to respond as quickly as they could by pressing the space bar on a computer keyboard. During stop trials, the imperative signal was followed after a stop-signal delay (SSD) by the stop signal, which was a red circle. Participants had to refrain from responding, i.e., cancel and inhibit their response to the imperative signal, upon presentation of the stop signal.

The majority of trials (70% of total trials) were ‘go’ trials, whereby the imperative signal was presented for a maximum of 1500 ms, or until a response was provided. This was followed by the display of feedback for 1000 ms, where the participant’s go reaction time (go RT: the duration from imperative signal onset to response) was displayed. If no response was provided within the 1500-ms time limit, ‘missed’ was displayed on the screen.

During stop trials (30% of total trials; randomly interspersed in each block), the imperative signal was replaced by the stop signal, which was a red circle, after a stop-signal delay (SSD). The initial SSD was set at 50 ms. During stop trials, participants had to attempt to cancel (inhibit) the go response and *not* press the space bar. Stop signals were displayed for 1000 ms, or until a response (a failed stop) occurred. The trial feedback – ‘good!’ for successful stops and ‘wrong response’ for failed stops – was then displayed for 1000 ms.

The SSD was adjusted in a staircase fashion, wherein a successful stop would result in an increase in SSD of 50 ms for the following stop trial, while a failed (unsuccessful) stop would result in SSD being reduced by 50 ms for the subsequent stop trial. The minimum SSD was 50 ms. This tracking procedure aims for a 0.5 probability of a failed stop trial (P(Respond|Stop) ∼ 0.5), thus improving the reliability of SSRT estimates (Band et al., 2003; Verbruggen et al. 2019).

Participants performed a practice block of 20 trials (6 stop trials) before the first pre-tACS behavioral assessment to ensure that they understood and adhered to task instructions (Verbruggen et al., 2019). The practice block was administered a second time for individuals who might not have understood or adhered to the task instructions during the first practice block (e.g., if a participant appeared to have delayed their response to the imperative signal in anticipation of the stop signal). Two blocks of the stop-signal task were administered at each assessment time (pre-tACS and post-tACS). Each assessment block contained 100 trials (block duration: ∼4 min; 30 stop trials per block).

### 2.4. tACS

High-definition tACS was delivered double-blind, using a neuroConn DC-Stimulator MC (neuroCare Group GmbH). Rubber disc electrodes of 2 cm diameter were placed on the scalp using Ten20 conductive paste in a 2 X 1 montage for each stimulation site (see Figure 3). The rIFG was localized at the intersection point of the line between T4 and Fz, and the line between F8 and Cz (10-10 electrode positions; Chatrian et al., 1985; Jacobson et al., 2011). The preSMA was localized at Fz (Hsu et al., 2011). Electrode impedance was kept below 50 kΩ. Sinusoidal currents at 20 Hz (beta frequency) with zero DC offset were applied for a stimulation duration of 20 minutes, with a 30 s ramp-up/down of currents at the beginning and end of stimulation.

**Figure 3.**
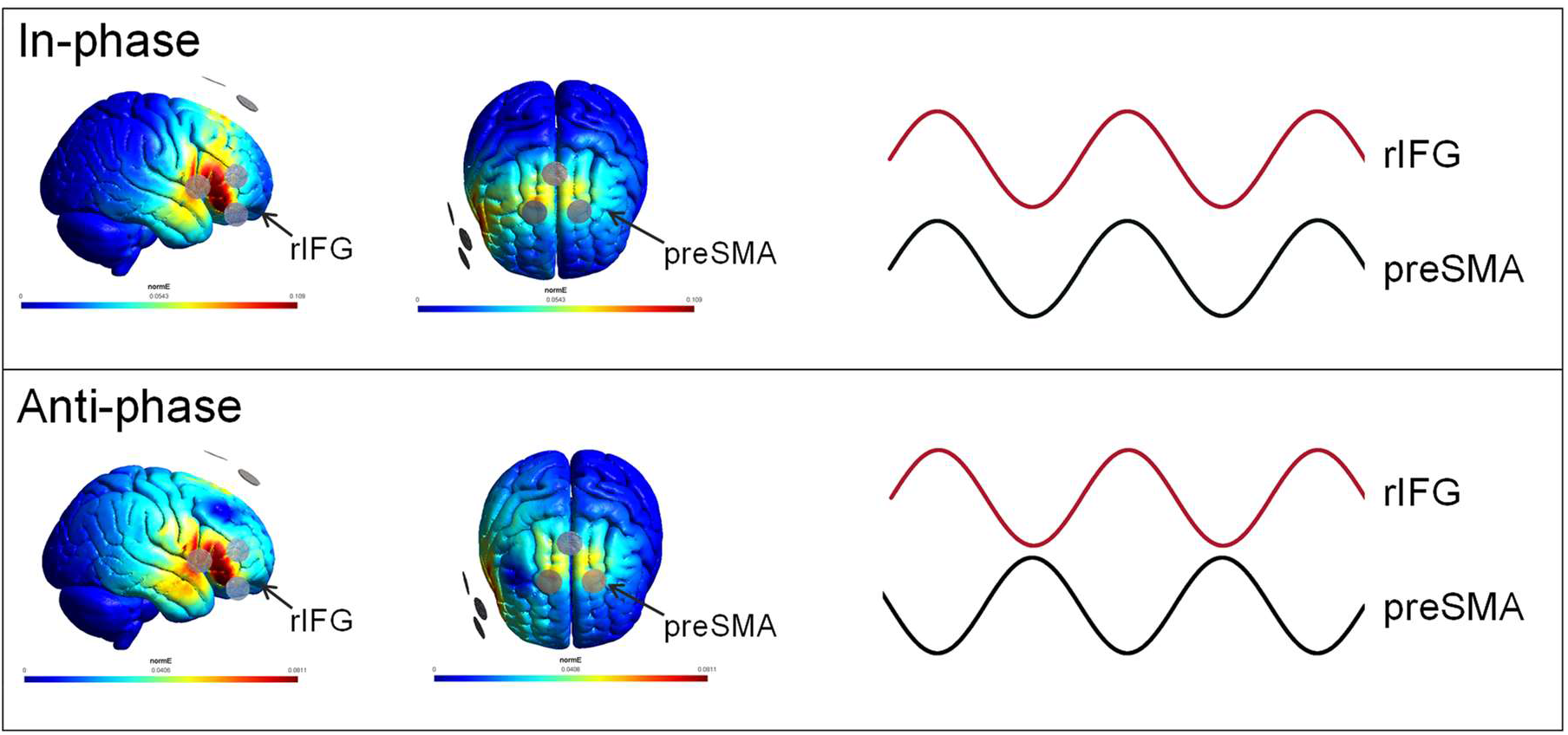
Current Flow Model. *Note.* A 2 X 1 electrode montage was used for each stimulation site. Currents with peak-to-peak amplitudes of 1 mA were delivered to the stimulation sites with a 0° and 180° phase difference for in- and anti-phase tACS, respectively.

The electric field model is depicted in Figure 3. The currents were applied with peak-to-peak amplitudes of 1 mA to both stimulation sites at 0° phase lag and 180° phase lag for in-phase and anti-phase tACS conditions, respectively.

### 2.5. EEG Acquisition and Preprocessing

EEG was recorded using a Net Amps 300 amplifier and Net Station (4.5.6) software, with 128-channel HydroCel Geodesic Sensor Nets (GSN, Magstim EGI; egi.com) and the GSN 128 1.0 montage. Data were acquired at a sampling rate of 1000 Hz, and online referenced to the vertex (Cz). Impedance was kept below 50 kΩ. The EEG signals were high-pass filtered at 0.1 Hz and low-pass filtered at 500 Hz during recording.

The EEG data were preprocessed with the EEGLAB toolbox (Delorme & Makeig, 2004) through the MATLAB environment (MathWorks, R2020a). The data were down-sampled to 500 Hz, bandpass filtered from 1 to 85 Hz, and notch filtered at 50 Hz. Resting-state data were divided into 2 s epochs. Task-related data were epoched into time windows that were time-locked to the stop signal (-500 to 1200 ms) and baseline-corrected using prestimulus activity from -500 to 0 ms. The epoched data were visually inspected, with bad channels and noisy epochs manually removed. Following the interpolation of removed channels, the *fullRankAveRef* EEGLAB plugin was used to re-reference the data to the average. Finally, an independent component analysis (Informax algorithm) was conducted and the components were visually inspected for the visual detection and removal of ocular, vascular, and myogenic artifacts.

A surface Laplacian spatial filter was applied to both task-related and resting-state data. This filter improves the spatial resolution of scalp-level activity by attenuating spatially-broad activity that is likely due to volume conduction (Nunez & Srinivasan, 2006). The surface Laplacian was computed using Perrin et al.’s (1989) spherical spline method (smoothing parameter of 10^-5^ and Legendre polynomial order of 40) with the MATLAB implementation by Cohen (2014).

### 2.6. Data Analyses

#### 2.6.1. Stop-Signal Task Performance

SSRTs were estimated using the integration method, which involves the subtraction of the mean SSD from the reaction time at which the integral of the go RT (reaction time of go trials) distribution equals zero (Logan & Cowan, 1984). While the integration method does not require that P(Respond|Stop) = 0.5, probabilities that are close to 0.5 improve the reliability of SSRT estimates (Band et al., 2003). For that reason, the data from participants with P(Respond|Stop) lower than 0.25 or higher than 0.75 were excluded from all analyses (Congdon et al., 2012; Verbruggen et al., 2019). We assessed this via stop accuracy, the inverse construct of P(Respond|Stop), i.e., the proportion of correctly inhibited trials.

SSRTs and go RTs were analyzed using mixed effect models with fixed effects of AGE (young, older), TIME (pre, post), and STIM (in-phase, anti-phase) with the maximal random effect structure (i.e., by-participant random intercepts and by-participant random slopes for all fixed effects as well as correlations among random effects) as allowed by the data and justified by the design used for every model (Barr et al., 2013; Singmann & Kellen, 2019). SSRTs were analyzed with a linear mixed effect model (LMM), using mean SSRTs for the pre and post time points of each session (as SSRTs cannot be calculated at the single trial level). As the go RT data were not normally distributed, go RT data were analyzed using a generalized linear mixed effect model (GLMM) applying a gamma distribution with a log link function (Lo & Andrews, 2015). Go RT data at the single trial level was used for the GLMM. Null hypothesis significance testing for main and interaction effects was conducted using Wald Chi-Squared tests for GLMM and *F*-tests for LMM. As the effects of tACS on stop-signal task performance were of primary interest in the present study, only main effects and interactions involving TIME and STIM as factors will be described in detail. Significant effects were further investigated with Bonferroni-corrected contrasts. Effect sizes are reported as Cohen’s *d*, with *d* ≤ 0.2 indicating a small effect, 0.2 < *d* ≤ 0.5 for a medium effect, and *d* ≥ 0.8 for a large effect (J. Cohen, 1988). Statistical significance was set at *α* = .05.

#### 2.6.2. EEG Analyses

##### 2.6.2.1. Source Reconstruction

The localization of source activity was performed using depth-weighted linear L2- minimum norm estimates (Baillet et al., 2001; Gramfort et al., 2014) using the Brainstorm toolbox (Tadel et al., 2011). The forward model was constructed with the ICBM152 template brain (Fonov et al., 2009) and the Symmetric Boundry Element Method (BEM) implemented in the OpenMEEG software (Gramfort et al., 2010; Kybic et al., 2005). Dipole orientations for the source model were constrained to be orthogonal to the cortex.

The Desikan-Killiany Atlas (Desikan et al., 2006) was used for cortical parcellation of the right IFG into its components: the pars opercularis, pars orbitalis, and pars triangularis. The centre coordinates of the preSMA region (left preSMA; area = 28 vertices/ 6.52 cm^2^) were based on the regional activations recorded during stop-signal inhibition by Li et al. (2006): x = -4, y = 36, z = 56 (conversion from Talairach to MNI coordinates by Duann et al., 2009). Due to the placement of the tACS electrodes that were targeted at preSMA, any tACS-induced effects would likely be distributed to both left and right preSMA (see current flow model in Figure 3). As such, the homologous area in the right hemisphere (right preSMA; MNI coordinates (mm): x = 4, y = 36, z = 56; area = 28 vertices/ 6.15 cm^2^) was also included in the source-level analysis.

We constrained our source-level analyses to cortical regions as the accuracy of subcortical source estimates is a subject of debate (Attal & Schwartz, 2013; Krishnaswamy et al., 2017). This is due in part to the depth and cytoarchitecture of subcortical structures, which makes it difficult to detect their activity using EEG (Attal & Schwartz, 2013), and also due to the challenges in delineating subcortical from cortical activity (Krishnaswamy et al., 2017). Despite evidence suggesting that the localization of subcortical sources is feasible (Krishnaswamy et al., 2017; Seeber et al., 2019), the accuracy of their source estimates is still considerably lower than for the neocortex, particularly for thalamic regions (Attal & Schwartz, 2013). The separation of subcortical and cortical activity is also compromised if the latter is not confined to a subset of cortical regions (Krishnaswamy et al., 2017).

##### 2.6.2.2. Imaginary Component of Coherency (ImCoh)

The analyses of rIFG-preSMA connectivity were performed based on source localized signals from bilateral preSMA, and the right pars opercularis, pars orbitalis, and pars triangularis (components of the rIFG). The signals were convolved with complex Morlet wavelets in 1-Hz increments for frequencies between 8 and 45 Hz. The wavelet lengths ranged from 4 to 10 cycles in 38 linearly-spaced steps, with the wavelet length increasing with the wavelet frequency for the dynamic adjustment of the balance between temporal and frequency precision (M. X. Cohen, 2014). To minimize the effects of edge artifacts, the resultant analytic signals were analyzed in time windows of 400 to 1600 ms in each epoch for resting state data and -200 to 800 ms for task-related data.

The imaginary component of coherency (ImCoh), an index of the consistency of phase angle differences (phase lag) between signals (Nolte et al., 2004), was used to assess changes in phase-coupling that were induced by tACS. For channels *i* and *j*, with the complex Fourier transforms *x_i_(f)* and *x_j_(f)* of their time series data, ImCoh at frequency *f* is given as

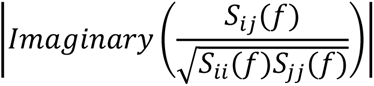

where the cross-spectral density *S_ij_(f)* is derived from the complex conjugation of *x_i_(f)* and *x_j_(f)*. The coherency between the channel signals is obtained by normalising the cross-spectral density by the square root of the signals’ spectral power (*S_ii_(f)* and *S_jj_(f)*). The result is a complex number from which the imaginary component, ImCoh, is extracted. ImCoh has been shown to be insensitive to the effects of volume conduction, and its usage thus reduces the likelihood of mistakenly taking spurious connectivity for real functional coupling (Nolte et al., 2004). An ImCoh value of 1 denotes perfect phase-coupling between signals, while a value of zero indicates that the phase angle differences between the signals are completely random. The ImCoh was calculated for each inter-region pair (left preSMA-pars opercularis, left preSMA-pars orbitalis, left preSMA-pars triangularis, right preSMA-pars opercularis, right preSMA-pars orbitalis, and right preSMA-pars triangularis) and subsequently averaged across all pairs to result in the mean ImCoh estimate between the rIFG and preSMA (rIFG-preSMA ImCoh for brevity).

The estimates of ImCoh were computed at the trial-level, in sliding time windows with frequency-varying lengths of 3-8 cycles in 38 linearly-spaced steps, at 20-ms intervals within each epoch. For resting-state data, these estimates were subsequently averaged within and across epochs to obtain an ImCoh estimate for every participant at pre- and post-tACS for in- and anti-phase tACS sessions. The ImCoh estimates for task-related data were baseline-corrected to -200-0 ms from stop-signal onset and averaged across epochs at each time point. This was conducted for the stop trials of every participant at pre- and post-tACS for in- and anti-phase tACS sessions.

To determine the frequency-specificity of tACS-induced effects on ImCoh, we assessed the changes in rIFG-preSMA ImCoh from pre-to post-tACS for each age group and tACS condition. This analysis was performed with ImCoh estimates from resting-state data and from successful stop trials for task-related data. Through cluster-based permutation testing, the ImCoh values at pre- and post-tACS were contrasted at each sample, i.e., frequency point for resting-state data and time-frequency point for task-related data. This non-parametric statistical approach allows multiple univariate tests to be carried out for each sample while controlling for the family-wise error rate (Maris, 2012; Maris et al., 2007). The permutation distributions of maximum cluster sizes were constructed with 2000 permutations. In each permutation, the session time labels (i.e., pre-tACS and post-tACS) were randomly assigned to the ImCoh values at each sample. A paired-sample *t*-test was performed to compare the ImCoh values between the randomly assigned time labels at each sample.

The correlation between the change from pre-to post-tACS for ImCoh (ΔImCoh = ImCoh_post-tACS_ - ImCoh_pre-tACS_) and response inhibition performance (ΔSSRT = SSRT_post-tACS_ - SSRT_pre-tACS_) was also analyzed with cluster-based statistics. The ΔSSRT and ΔImCoh at each sample point were ranked across participants. The ranking values were subsequently used for the computation of the Spearman’s correlation coefficient (*ρ*) between SSRTs and sample-level ImCoh values. During each permutation of the cluster-based analyses, the ranking values for ΔSSRT were randomly shuffled at each sample across participants before *ρ* was computed. For the thresholding of sample points, *ρ* values were converted into *t* values using the formula:

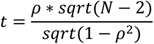

where *N* was the number of participants in the subgroup that was being analyzed.

The cluster-based permutation tests were performed with *α* = .05 for the thresholding of statistical test values (e.g., *t* values) at each sample point. Sample points with suprathreshold values were clustered according to spectral adjacency for resting-state data, and temporal and spectral adjacency for task-related data, with separate clusters for positive and negative values. The size of each cluster was obtained by taking the sum of the absolute statistical values within the cluster. The permutation distribution was constructed with the largest cluster size of every permutation, and its 97.5th percentile was set as the cluster-level correction threshold. The clusters in the real (i.e., not permuted) data with cluster sizes that exceeded this threshold were deemed to be significant.

##### 2.6.2.3. Spectral Power Analyses

Changes in EEG coherency (and its imaginary component ImCoh) can also be due to changes in spectral power instead of real changes in phase-coupling (Nolte et al., 2004). To determine if the results of our ImCoh analyses might be confounded by significant changes in oscillatory power, we also examined the effects of tACS on spectral power during resting-state and successful stop trials.

The spectral power at each sample point was obtained by taking the square of the analytic signals that were derived from wavelet convolution. The power estimates were averaged across trials for task-related data, and across trials and within trials for resting-state data, before being converted to decibels (dB; 10log10(power)). Decibel conversion ensures that data across conditions and subjects are on the same scale, thus facilitating the comparison of power values (M. X. Cohen & van Gaal, 2013). Task-related changes in power estimates for task data (event-related spectral perturbation) were normalized to a baseline window of -200 to 0 ms pre-stop-signal onset before decibel conversion.

The changes in spectral power from pre-to-post tACS were assessed with cluster-based permutation statistics for rIFG and preSMA. For each region, the spectral power for pre- and post-tACS were contrasted at every sample point. To obtain the distributions of maximum cluster sizes, ‘Pre’ and ‘Post’ labels were randomly allocated to the power values at each sample point during each permutation.

Cluster-based statistics were also used to investigate the association between the tACS-induced changes in response inhibition (ΔSSRT) and the changes in spectral power (ΔPower = Power_post-tACS_ – Power_pre-tACS_) for preSMA and rIFG. Spearman’s *ρ* was computed with the ranking values of ΔSSRT and ΔPower for each sample point. In each permutation, Spearman’s *ρ* was computed for each sample point after the ranks for ΔSSRT were randomly shuffled across participants.

##### 2.6.2.4. Successful Versus Failed Stop Trials

The main premise of our hypothesis is that the functional connectivity between rIFG and preSMA underlies successful response inhibition. To verify that this is indeed the case for our study, we investigated if rIFG-preSMA ImCoh would be stronger during successful versus failed stop trials. For each age group (young and older), stimulation condition (in- and anti-phase tACS), and session time (pre- and post-tACS), rIFG-preSMA ImCoh was computed for every time-frequency point within successful and failed stop trials. The mean ImCoh during successful stop trials was subsequently obtained by averaging the ImCoh values across time points in the interval between stop-signal onset (0 ms) and the SSRT of each participant. This provided estimates of phase connectivity that preceded successful inhibition – i.e., before the SSRT, the time at which the internal response to the stop signal occurs (Logan & Cowan, 1984). Conversely, the phase connectivity preceding stop failures was obtained by averaging the ImCoh values of failed stop trials across the time points that lie between stop-signal onset (0 ms) and the stop reaction time (reaction time of response due to failed response inhibition) of each participant.

Cluster-based permutation tests were used to investigate the difference in ImCoh between successful and failed stop trials. A paired-samples *t-*test was used to contrast the ImCoh values between the two trial types at every frequency point. During each permutation, the ImCoh values at each frequency point were randomly assigned as belonging to a ‘successful’ or ‘failed’ stop trial.

Unless otherwise stated, statistical analyses and visual illustration of the results were performed via customized scripts in MATLAB (MathWorks, R2020a), and the software package R for Statistical Computing version 3.6 (R Core Team, 2019) using the packages ‘tidyverse’ (Wickham, 2021), ‘ggpubr’ (Kassambara, 2020), ‘DescTools’ (Signorell, 2021), ‘janitor’ (Firke, 2021), ‘reshape2’ (Wickham, 2020), ‘rcompanion’ (Mangiafico, 2021), ‘car’ (Fox et al., 2020), ‘heplots’(Fox & Friendly, 2021), ‘rstatix’ (Kassambara, 2021), ‘Hmisc’ (Harrell, 2021), ‘rColorBrewer’ (Neuwirth, 2014), and ‘hrbrthemes’ (Rudis, 2020).

#### 2.6.3. Data and Code Availability Statement

Data are available upon request to the authors, subject to approval from Murdoch University Human Research Ethics Committee. Analysis codes are publicly available on the Open Science Framework (OSF): https://osf.io/mh9ks/.

## 3. Results

Unless otherwise stated, data are expressed as mean (*M*) ± standard deviation (*SD*), and statistical significance was based on an alpha level of 0.05.

### 3.1. Control Measures

There were no significant differences in perceived tACS sensations, sleep quality, sleep quantity, caffeine intake, and alcohol intake between the different tACS conditions (see Supplementary Material Tables S1-S2). This suggests that our findings were unlikely to have been significantly impacted by these potential confounds.

### 3.2. Stop-Signal Task Performance

The descriptive statistics of SSRT, go RT, failed stop RT (reaction time on failed stop trials), stop accuracy, and go accuracy are presented in Table 1.

**Table 1.** Stop-Signal Task Performance

| Outcome variable | Older |  |  |  | Young |  |  |  |
| --- | --- | --- | --- | --- | --- | --- | --- | --- |
|  | In-Phase tACS |  | Anti-Phase tACS |  | In-Phase tACS |  | Anti-Phase tACS |  |
|  | Pre | Post | Pre | Post | Pre | Post | Pre | Post |
| SSRT (ms) | 263 ± 32 | 256 ± 29 | 275 ± 48 | 262 ± 33 | 229 ± 28 | 218 ± 30 | 222 ± 37 | 215 ± 37 |
| Go RT (ms) | 494 ± 152 | 537 ± 148 | 500 ± 143 | 528 ± 153 | 332 ± 103 | 319 ± 101 | 335 ± 109 | 328 ± 110 |
| Failed Stop RT (ms) | 406 ± 108 | 451 ± 138 | 411 ± 107 | 435 ± 119 | 273 ± 76 | 270 ± 88 | 290 ± 110 | 281 ± 109 |
| Stop accuracy (%) | 57.67 ± 6.00 | 58.44 ± 4.98 | 56.22 ± 6.25 | 58.22 ± 6.91 | 47.87 ± 6.34 | 47.59 ± 5.75 | 48.33 ± 7.41 | 47.69 ± 8.42 |
| Go accuracy (%) | 99.38 ± 1.29 | 99.24 ± 0.95 | 99.38 ± 0.71 | 99.57 ± 0.45 | 99.84 ± 0.31 | 99.40 ± 1.67 | 99.68 ± 0.56 | 99.33 ± 1.16 |

*Note.* Data are expressed as mean ± standard deviation.

#### 3.2.1. Stop-Signal Reaction Time (SSRT)

As expected, there was a significant main effect of AGE, *F*(1, 35.13) = 19.01, *p* = 0.0001, with young participants (*M* = 220.97 ms, *SD* = 32.64 ms) exhibiting significantly shorter SSRTs (i.e., better response inhibition) when compared with older participants (*M* = 264.54 ms, *SD* = 36.01 ms) averaged across the factors TIME and STIM. There was also a significant main effect of TIME, *F*(1, 105.39) = 7.21, *p* = 0.008, indicating that SSRTs at post-tACS (*M* = 234.65 ms, *SD* = 38.22 ms) were significantly shorter than at pre-tACS (*M* = 245.24 ms, *SD* = 41.48 ms), irrespective of age group and tACS condition. There were no significant AGE X STIM X TIME and STIM X TIME interactions, *F*s < 2.80, *p*s > 0.097. This suggests that there were no significant changes in SSRTs that were phase-specific for young and older participants. See Figure 4 for the SSRTs of both age groups at each time point for both tACS conditions.

**Figure 4.**
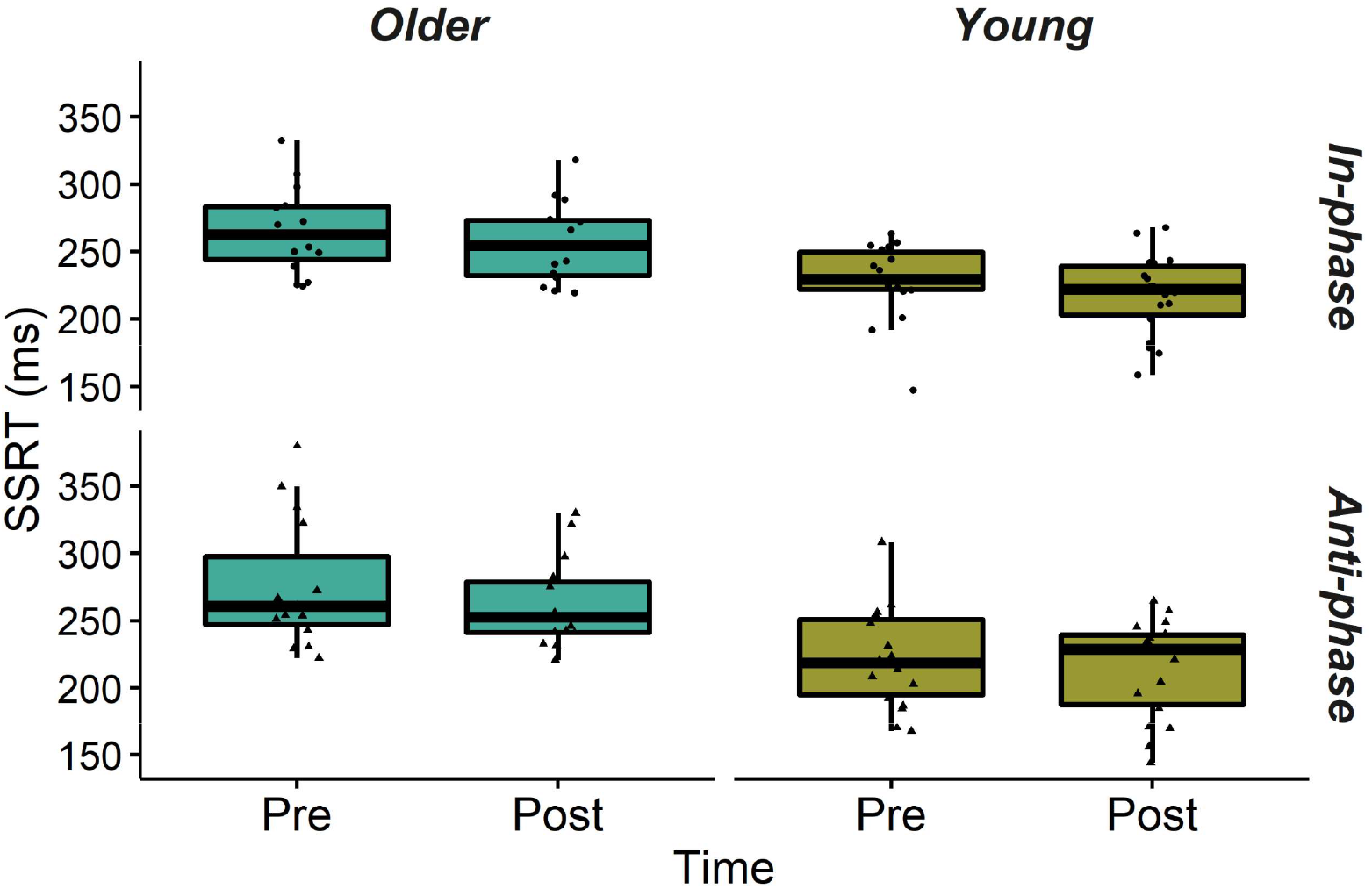
Stop-Signal Reaction Time (SSRT) *Note*. The LMM for SSRT indicate no significant STIM X TIME interactions, suggesting that improvements in SSRTs did not vary according to in- or anti-phase tACS for either age group.

#### 3.2.2. Go Reaction Time (Go RT)

The main effect of AGE, χ*^2^*(1, *N* = 33) = 524.34, *p* < 0.0001, and interaction between AGE and TIME, χ*^2^*(1, *N* = 33) = 25.75, *p* < 0.0001, were significant for go RT. These were mediated by a higher order three-way interaction between AGE, TIME, and STIM, χ*^2^*(1, *N* = 33) = 4.43, *p* = 0.035. Post-hoc analyses found that older participants had significantly longer go RTs after both in-phase tACS, *z* = 4.60, *p* < 0.0001, *d* = 0.34, and anti-phase tACS, *z* = 2.86, *p* = 0.004, *d* = 0.19, while young participants had significantly shorter go RTs after in-phase tACS, *z* = -2.59, *p* = 0.0097, *d* = -0.18. There were no significant changes in go RTs for young participants after anti-phase tACS, *z* = -1.41, *p* =0.158, *d* = -0.09 (see Figure 5).

**Figure 5.**
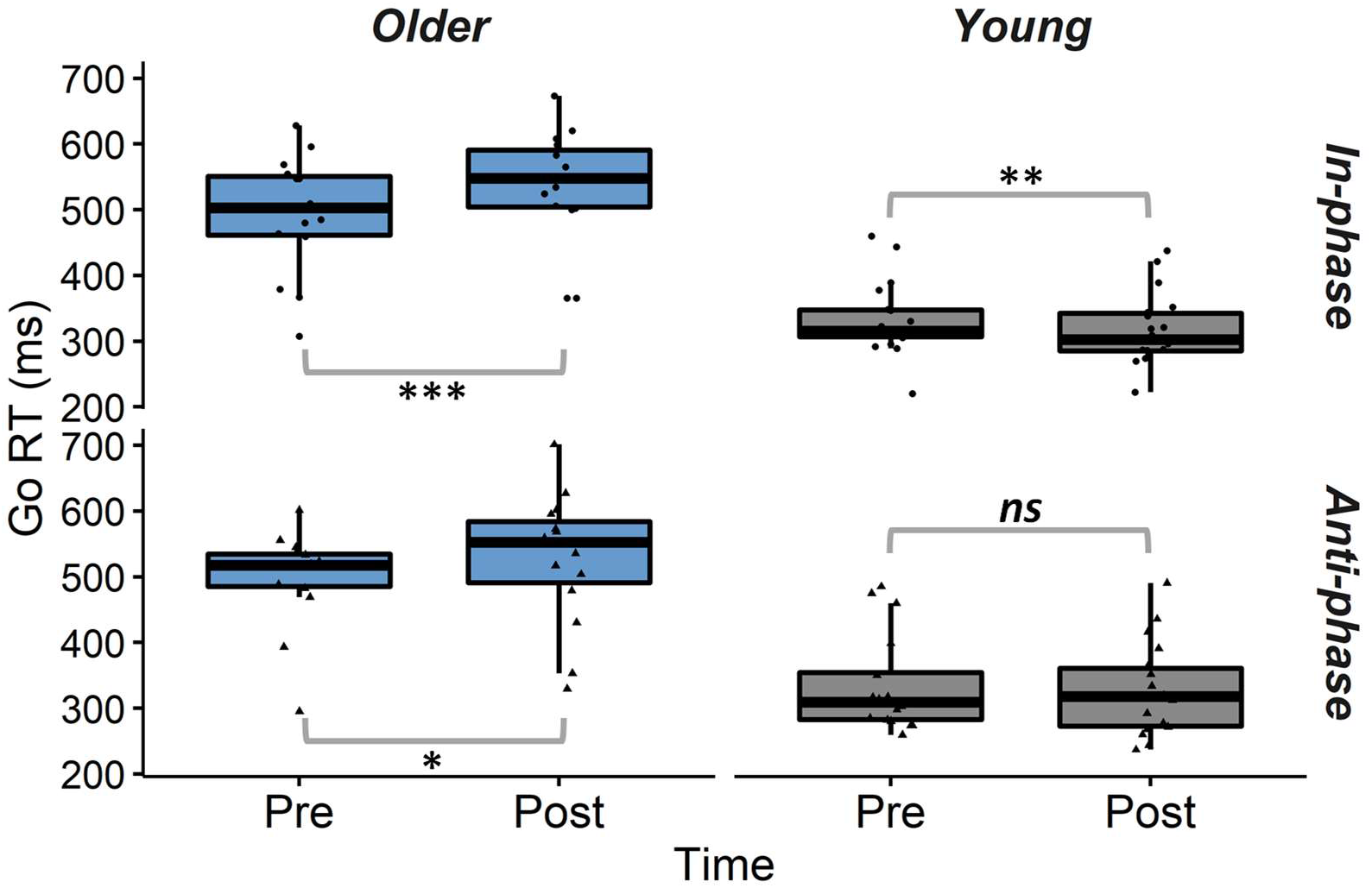
Go Reaction Time (Go RT) *Note*. Older participants exhibited significantly longer go RTs after in- and anti-phase tACS, while young participants had significantly shorter go RTs after in-phase tACS. There were no significant changes in go RTs for young participants after anti-phase tACS. *ns*: not significant * *p* < 0.01, ** *p* < 0.001, *** *p* < 0.0001.

### 3.3. rIFG-preSMA ImCoh

Cluster-based permutation testing found no significant differences between the pre- and post-tACS estimates of resting-state and task-related ImCoh for in-phase or anti-phase tACS in young and older participants. There were also no significant correlations between ΔImCoh and ΔSSRT after in- and anti-phase tACS for young and older participants (see Supplementary Material Figures S1-S4).

### 3.4. Spectral Power Analyses

There was a significant increase in resting-state spectral power (11-45 Hz) at the rIFG for young adults after anti-phase tACS (see Figure 6). There were no other significant pre-post changes in resting-state (Figure 6) and task-related (Supplementary Material Figure S5) spectral power at the rIFG and preSMA for either age groups from in-or anti-phase tACS.

**Figure 6.**
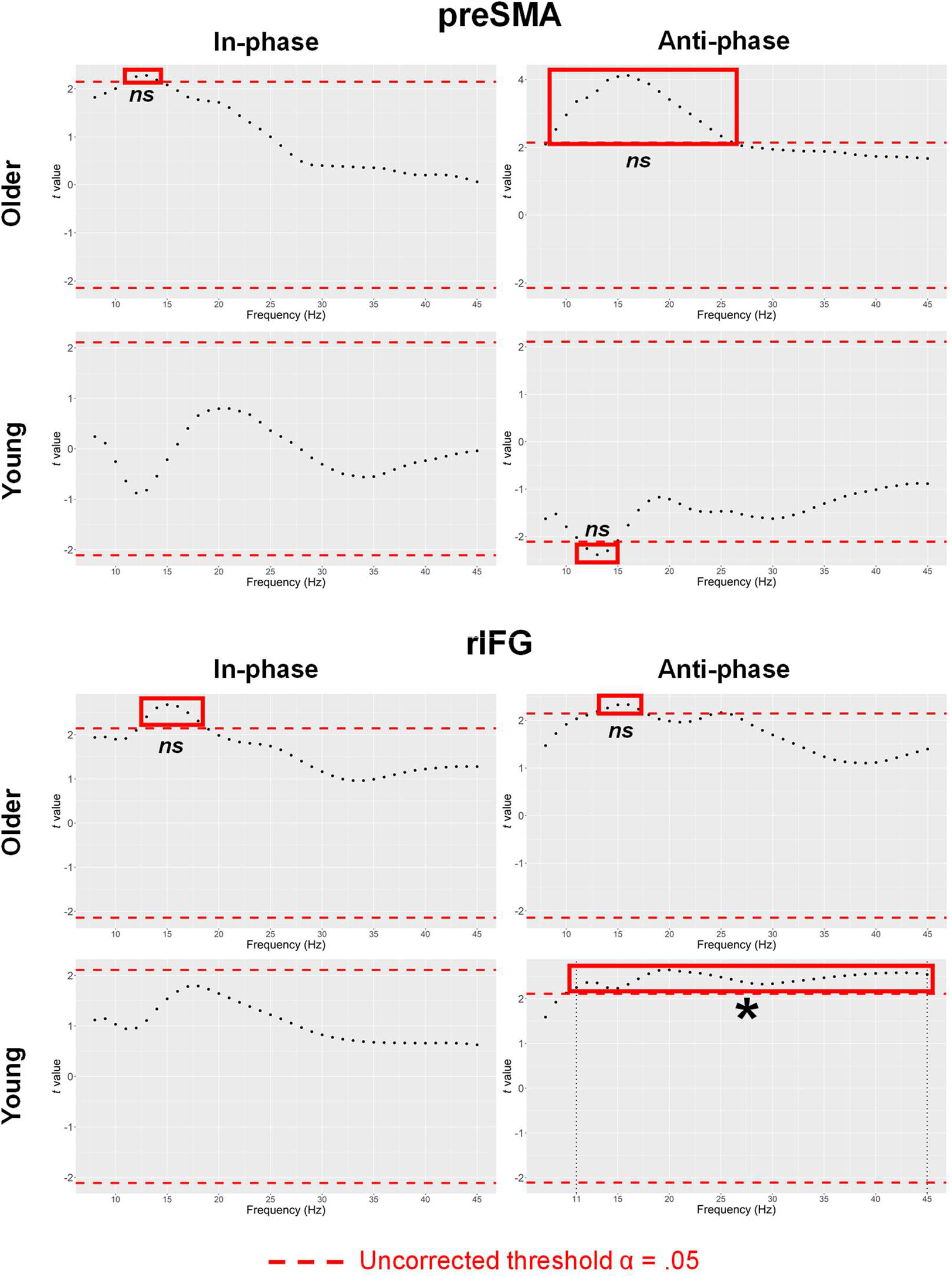
Cluster-Based Analyses of Pre-to Post-tACS Changes in Resting-State Spectral Power at preSMA and rIFG. *Note*. The *t* values for the pre-to post-tACS difference in power spectral values at each spectral point. The red dashed lines represent the uncorrected threshold, i.e., critical *t* at α = .05 before cluster-based correction. There was a significant increase in rIFG power from 11 to 45 Hz for young participants after receiving anti-phase tACS. There were no other significant changes in pre-post spectral power for rIFG and preSMA in either young or older participants in any of the other conditions. *ns*: not significant after cluster-based correction. *significant after cluster-based correction.

The analyses of correlations between ΔSSRT and ΔPower found a significant negative correlation between task-related beta power (∼14-33 Hz) and SSRTs for older participants after receiving in-phase tACS, with stronger beta power at the preSMA around the 0-200ms time-window of successful stop trials associated with shorter SSRTs (i.e., better response inhibition) (see Figure 7). There were no significant correlations between ΔSSRT and ΔPower at resting-state (see Supplementary Material Figure S6).

**Figure 7.**
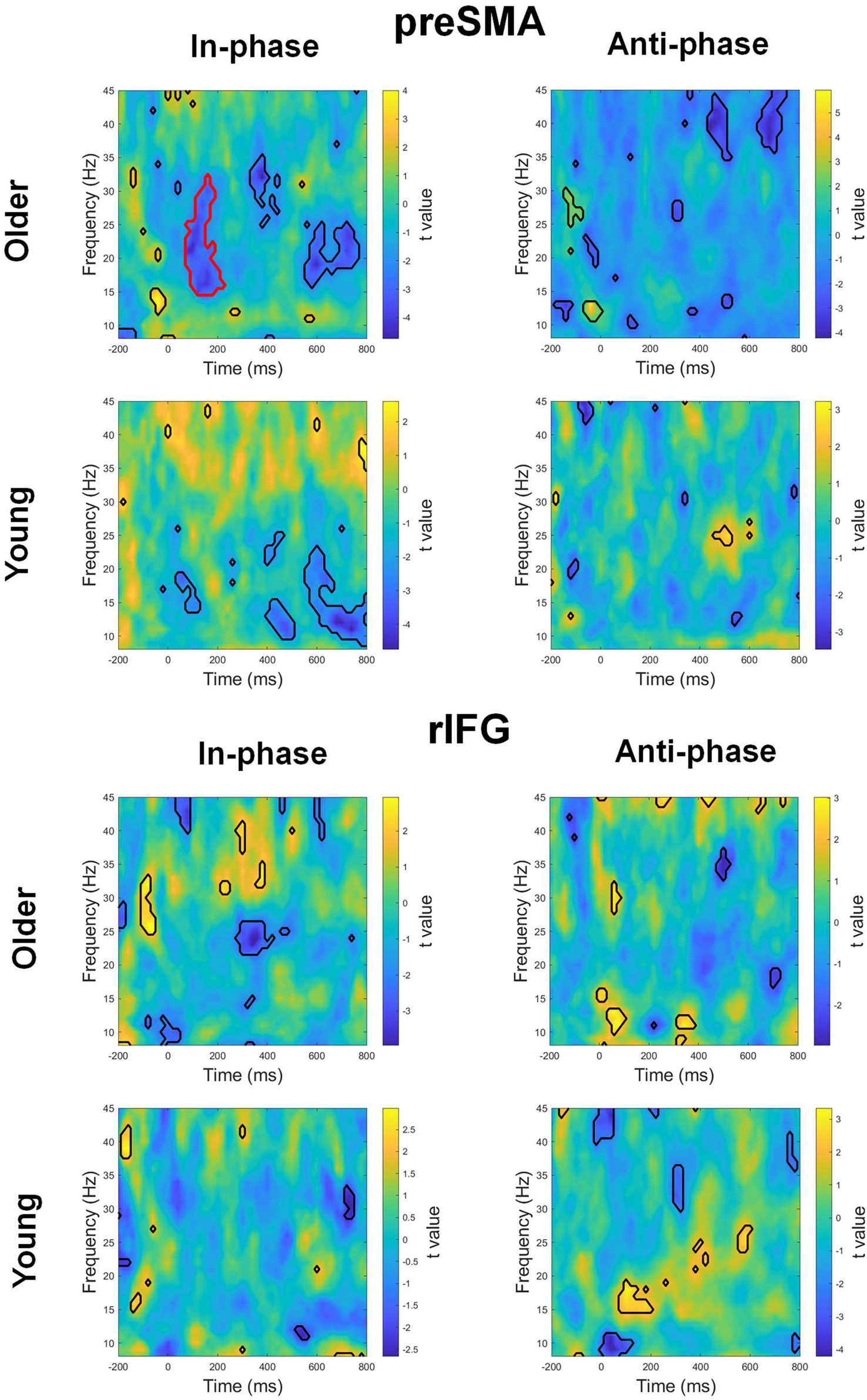
Cluster-Based Analyses of Correlation between ΔSSRT and ΔPower at preSMA and rIFG During Successful Stop Trials. *Note*. Time-frequency plots showing the *t* values (converted from Spearman’s *ρ*) for the correlation between ΔPower and ΔSSRT at each time-frequency point during successful stop trials. Regions of suprathreshold clusters are denoted by black outlines (*p* < .05, uncorrected). A cluster in the in-phase tACS condition for older participants (outlined in red) was found to be significant after cluster-based correction, indicating that increases in beta power at the preSMA was

### 3.5. Successful Versus Failed Stop Trials

There were no significant differences in ImCoh between successful and failed stop trials for young and older participants before and after in- and anti-phase tACS (see Supplementary Material Figure S7).

## 4. Discussion

Our primary hypothesis was that the delivery of in-phase tACS at beta frequency (20 Hz) would strengthen phase connectivity between the stimulated regions and improve response inhibition in healthy young and older adults. Conversely, anti-phase tACS was expected to weaken phase connectivity and degrade response inhibition.

Contrary to our hypotheses, there were no phase-specific changes in response inhibition performance and rIFG-preSMA connectivity. Response inhibition was significantly improved (significantly shorter SSRTs) from pre-to post-tACS, but this improvement occurred across young and older participants, and across tACS conditions (in- and anti-phase). This indicates that the changes in SSRTs were not due to any phase-specific effects of our tACS protocol. It is possible that these improvements in SSRTs were instead the result of tACS-induced changes plasticity – beta tACS has previously been shown to induce NMDAR-mediated plasticity in the primary motor cortex (Wischnewski, Engelhardt, et al., 2019).

Further research is required to investigate the potential neuroplastic effects of this dual-site tACS protocol and their effects on response inhibition.

There were also no significant tACS-induced changes in functional connectivity between the preSMA and rIFG, which suggests that the dual-site tACS protocol was not effective in modulating rIFG-preSMA connectivity. Notably, our exploratory analysis also found that rIFG-preSMA connectivity was not significantly different between successful and failed stop trials. This indicates that rIFG-preSMA connectivity – specifically, phase connectivity as operationalized by ImCoh – might not underlie response inhibition. This is surprising, considering that a number of studies have implicated functional connections between the rIFG and preSMA in successful response inhibition (Dambacher et al., 2014; Duann et al., 2009; Jahfari et al., 2011, 2012; Rae et al., 2015). However, given that these studies measured functional connectivity from blood oxygen-level dependent signals, it is possible that our contrary findings were due to our use of electrophysiological signals (i.e., EEG) for the determination of functional connectivity. This warrants future investigation into the potential differences in how hemodynamic and electrophysiological connectivity underlie cognitive processes.

Interestingly, our spectral power analyses found that increases in beta power at the preSMA were significantly associated with better response inhibition (shorter SSRTs) for older participants after in-phase tACS. This corroborates the findings of a recent study, in which 20 Hz beta tACS over the preSMA enhanced motor inhibition during stop-signal task performance (Leunissen et al., 2022). Participants in that study responded to the go stimulus by pressing a force sensor, and event-related tACS was pseudo-randomly applied on 40% of the stop-signal trials. The study found that 20 Hz tACS applied over the preSMA significantly decreased peak force and peak force rate on successful stop trials. Taken together, this suggests that 20 Hz beta tACS over the preSMA might be efficacious in improving response inhibition. The lack of significant correlation between ΔSSRT and ΔPower for the young participants in our study could be due to potential ceiling effects in response inhibition performance. It is also possible that more subtle changes in response inhibition were not captured, since the stop-signal task measures motor inhibition in a dichotomic manner i.e., stop trials are either successful or unsuccessful (Nguyen et al., 2021). For instance, changes in “partial” responses could not be ascertained from SSRTs (Nguyen et al., 2021). The inclusion of a force sensor component (as used by Leunissen et al., 2022) for the force measurement of partial responses might have yielded a more sensitive measure of motor inhibition in our study, particularly if ceiling effects were present in the young participants’ stop-signal task performance.

Contrary to Leunissen et al. (2022), who found that 20 Hz tACS over preSMA led to significantly higher beta power around stop signal onset during successful stop trials when comparing between stimulated and unstimulated trials, we found no significant changes in task-related spectral power from our tACS protocol, in which 20 Hz tACS was also applied over preSMA. This might be due to differences in stimulation montage, with our 2 X 1 montage potentially resulting in less focal electric fields when compared with Leunissen et al.’s (2022) 4 X 1 montage. Furthermore, current flow between the tACS electrodes at the rIFG and preSMA in our tACS protocol might have resulted in the co-stimulation of brain areas between the two regions and current shunting through the skin where the electrodes of each region were closest to each other (Saturnino et al., 2017) – this might have led to a less focal electric field at the preSMA, compared to a single-site protocol where only the preSMA is stimulated. However, it remains unclear as to why there was a significant decrease in resting-state spectral power (11-45 Hz; i.e., alpha to gamma frequencies) at the rIFG for young adults after anti-phase tACS. Speculatively, the application of anti-phase beta currents might have modulated cross-frequency phase-amplitude interactions (Canolty & Knight, 2010) that resulted in this increase in alpha to gamma power, but this requires future empirical investigation.

A key limitation of the current study was the disparity in tACS-induced field strengths at the rIFG and preSMA. Our current flow models (Figure 3) suggest that the tACS-induced electric field distribution at the preSMA was relatively weaker than that at rIFG, which might have limited the efficacy of the tACS protocol in modulating phase synchrony (Tan et al., 2020). It is also possible that the current intensity was too low for the significant modulation of functional connectivity – a review of tACS studies on corticospinal excitability found that intensities with peak-to-peak amplitudes above 1 mA elicited robust changes in corticospinal excitability, whereas intensities of 1 mA peak-to-peak resulted in less reliable changes in excitability (Wischnewski, Schutter, et al., 2019). Lastly, the lack of a sham condition in the current study also prevents a more conclusive investigation of the effects of our tACS protocol on response inhibition.

## 5. Conclusions

In- and anti-phase beta tACS over the rIFG and preSMA did not exert phase-specific effects on response inhibition, nor significantly modulated rIFG-preSMA connectivity for both young and older participants in this study. Interestingly, there were no significant changes in rIFG-preSMA phase connectivity between successful and failed stop trials, which suggests that rIFG-preSMA phase-coupling might not underlie effective response inhibition. Based on comparisons to other studies, our findings also suggest that beta tACS over the preSMA might have greater efficacy in improving response inhibition than a dual-site tACS protocol.

## CRediT author statement

**JT:** Conceptualization, Methodology, Formal analysis, Investigation, Software, Data Curation, Writing - Original Draft. **KI:** Software, Formal analysis, Writing - Review & Editing. **RP:** Formal analysis, Writing - Review & Editing. **MAN and MRH:** Writing - Review & Editing. **HF:** Conceptualization, Methodology, Formal analysis, Software, Data Curation, Writing - Review & Editing, Supervision.

## Funding

This work was supported by the Australia-Germany Joint Research Co-operation (DAAD) (57384703) awarded to HF, MAN, and JT, the Dementia Australia Research Foundation (711-1641) awarded to HF, MAN, and MRH. MRH was supported by the Australian Research Council Discovery Scheme (FT150100406; DP200101696).

### Declarations of interest

none

## Notes

### Competing Interest Statement

The authors have declared no competing interest.

### Summary of Updates

Stop-signal task performance was re-analysed with linear mixed models. Analyses of functional connectivity was performed on source-localized EEG data instead of sensor-level data.

