## Supplementary Material for "The Effects of Dual-Site Beta tACS over the rIFG and preSMA on Response Inhibition in Young and Older Adults"

### Control Measures

Planned comparisons using Wilcoxon signed-rank tests were performed to assess the perceived tACS sensations, sleep quality, sleep quantity, caffeine intake, and alcohol intake (see Table S1 for descriptive statistics) between the different tACS conditions for each age group. Effect sizes were measured with the Wilcoxon effect size, *r*, which was calculated with the R package rstatix’s *wilcox_effsize* function as the *z* statistic divided by square root of the sample size (*z*/√*n*), with *r* ≤ 0.1 for small effects, 0.1 < *r* ≤ 0.5 for medium effects, and *r* > 0.5 for large effects (Kassambara, 2021). There were no significant differences between the control measures between tACS conditions for either young or older participants (results of planned comparisons shown in Table S2).

Table S1

| Descriptive Statistics for Control Measures | | | | |
| --- | --- | --- | --- | --- |
|  | Older | | Young | |
| Control Measure | In-phase  tACS | Anti-phase  tACS | In-phase  tACS | Anti-phase  tACS |
| Sleep quality | 7.6 ± 1.35 | 7.6 ± 1.35 | 7.05 ± 1.48 | 6.78 ± 1.99 |
| Sleep quantity (hours) | 7.33 ± 0.98 | 7.27 ± 1.05 | 7.33 ± 1.60 | 7.19 ± 1.74 |
| Alcohol intake (units) | nil | nil | nil | nil |
| Caffeine intake (units) | 1.4 ± 0.83 | 1.47 ± 1.06 | 0.64 ± 0.83 | 0.75 ± 0.81 |
| Sensations during tACS | 2.66 ± 1.67 | 3.6 ± 3.33 | 4.66 ± 2.63 | 5.44 ± 7.59 |

*Note*. Data are expressed as mean ± standard deviation.

Table S2

| *Comparisons of Control Measures Between In- and Anti-Phase tACS Sessions* | | |
| --- | --- | --- |
| Control Measure | Older | Young |
| Sleep quality | *Z* = 38.00, *p* = .968,  *r* = 0.015 | *Z* = 40.50, *p* = .465,  *r* = 0.166 |
| Sleep quantity (hours) | *Z* = 20.50, *p* = .857,  *r* = 0.015 | *Z* = 57.00, *p* = .886,  *r* = 0.078 |
| Alcohol intake (units) | na | na |
| Caffeine intake (units) | *Z* = 4.00, *p* = .773,  *r* = 0.149 | *Z* = 15.00, *p* = .395,  *r* = 0.352 |
| Sensations during tACS | *Z* = 52.00, *p* = .325,  *r* = 0.281 | *Z* = 33.50, *p* = .418,  *r* = 0.203 |
| *Note*. Effect size is quantified by Cohen’s *r*, with *r* ≤ 0.1 for small effects, 0.1 < *r* ≤ 0.5 for medium effects, and *r* > 0.5 for large effects. na: not applicable. | | |

### rIFG-preSMA ImCoh

Cluster-based permutation testing found no significant differences between the pre- and post-tACS estimates of resting-state and task-related ImCoh for in-phase or anti-phase tACS in young and older participants. There were also no significant correlations between ∆ImCoh and ∆SSRT after in- and anti-phase tACS for young and older participants (Figures S1-4).

Figure S1

Cluster-Based Analyses of Pre- to Post-tACS Changes in Resting-State ImCoh

*Note*. The *t* values for the difference between pre and post ImCoh values at each spectral point. The red dashed lines represent the uncorrected threshold, i.e., critical *t* at α = .05 before cluster-based correction. There were no clusters of contiguous samples with suprathreshold *t* values.

Figure S2

Cluster-Based Analyses of Correlation Between ∆SSRT and ∆ImCoh at Resting State

*Note*. The *t* values (converted from Spearman’s *ρ*) for the correlation between ∆ImCoh and ∆SSRT at each spectral point. The red dashed lines represent the uncorrected threshold, i.e., critical *t* at α = .05 before cluster-based correction. There were no clusters of contiguous samples with suprathreshold *t* values.

Figure S3

Cluster-Based Analyses of Pre- to Post-tACS Changes in ImCoh During Successful Stop Trials

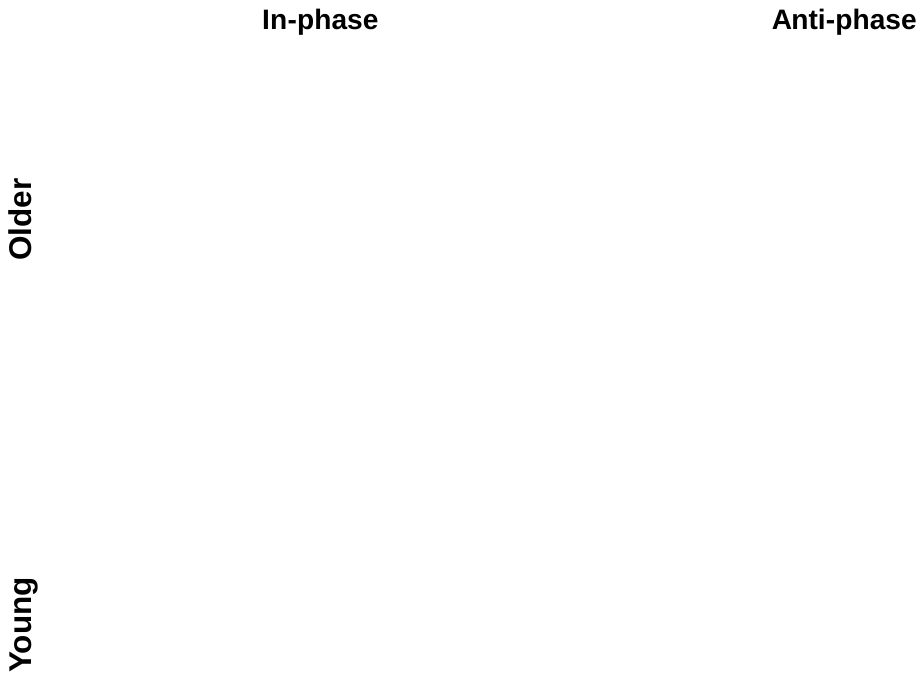

*Note.* Time-frequency plots showing the *t* values for all pre-tACS vs post-tACS contrasts of ImCoh during successful stop trials. Regions of suprathreshold clusters are denoted by black outlines (*p* < .05, uncorrected). None of the clusters survived cluster-based correction.

Figure S4

Cluster-Based Analyses of Correlation between ∆SSRT and ∆ImCoh During Successful Stop Trials

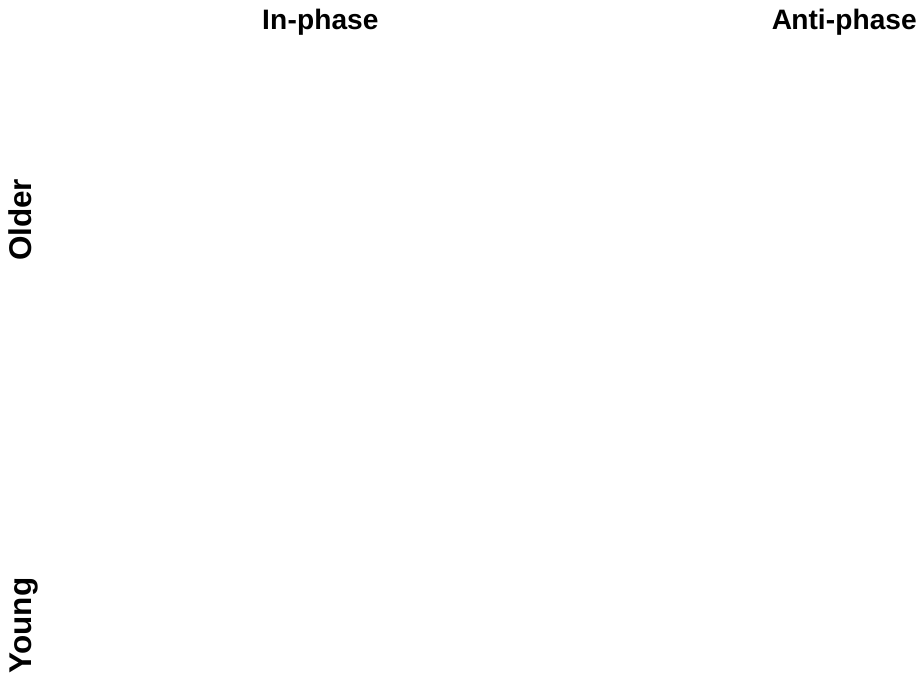

Note. Time-frequency plots showing the *t* values (converted from Spearman’s *ρ*) for the correlation between ∆SSRT and ∆ImCoh at each time-frequency point during successful stop trials. Regions of suprathreshold clusters are denoted by black outlines (*p* < .05, uncorrected). None of the clusters survived cluster-based correction.

### Spectral Power Analyses

There were no significant changes in task-related spectral power at the rIFG and preSMA for either age groups from in- or anti-phase tACS (Figure S5). There were no significant correlations between ∆SSRT and ∆Power at resting-state (Figure S6).

Figure S5

Cluster-Based Analyses of Pre- to Post-tACS Changes in Spectral Power at preSMA and rIFG During Successful Stop Trials

*Note.* Time-frequency plots showing the *t* values for all pre-tACS vs post-tACS contrasts of ImCoh during successful stop trials. Regions of suprathreshold clusters are denoted by black outlines (*p* < .05, uncorrected). None of the clusters survived cluster-based correction.

Figure S6

*Cluster-Based Analyses of Correlation Between ∆SSRT and ∆Power at preSMA and rIFG during Resting State*

*Note*. The *t* values for the correlation (converted from Spearman’s *ρ*) between ∆SSRT and ∆Power at each spectral point. The red dashed lines represent the uncorrected threshold, i.e., critical *t* at α = .05 before cluster-based correction. There were no clusters of contiguous samples with suprathreshold *t* values.

### Successful Versus Failed Stop Trials

There were no significant differences in ImCoh between successful and failed stop trials for young and older participants before and after in- and anti-phase tACS (Figure S7).

Figure S7

*Cluster-Based Analyses of Differences in ImCoh between Successful and Failed Stop Trials*

*Note*. The *t* values for the difference between pre and post ImCoh values at each spectral point. The red dashed lines represent the uncorrected threshold, i.e., critical *t* at α = .05 before cluster-based correction. *ns*: not significant after cluster-based correction.
